# Locating ligand binding sites in G-protein coupled receptors using combined information from docking and sequence conservation

**DOI:** 10.1101/461681

**Authors:** Ashley R. Vidad, Stephen Macaspac, Ho-Leung Ng

## Abstract

G-protein coupled receptors (GPCRs) are the largest protein family of drug targets. Detailed mechanisms of binding are unknown for many important GPCR-ligand pairs due to the difficulties of GPCR recombinant expression, biochemistry, and crystallography. We describe our new method, ConDock, for predicting ligand binding sites in GPCRs using combined information from surface conservation and docking starting from crystal structures or homology models. We demonstrate the effectiveness of ConDock on well-characterized GPCRs such as the β2 adrenergic and A2A adenosine receptors. We also demonstrate that ConDock successfully predicts ligand binding sites from high-quality homology models. Finally, we apply ConDock to predict ligand binding sites on a structurally uncharacterized GPCR, GPER. GPER is the G-protein coupled estrogen receptor, with four known ligands: estradiol, G1, G15, and tamoxifen. ConDock predicts that all four ligands bind to the same location on GPER, centered on L119, H307, and N310; this site is deeper in the receptor cleft than predicted by previous studies. We compare the sites predicted by ConDock and traditional methods that utilize information from surface geometry, surface conservation, and ligand chemical interactions. Incorporating sequence conservation information in ConDock overcomes errors introduced from physics-based scoring functions and homology modeling.

## Introduction

GPCRs (G-protein coupled receptors) are the largest family of drug targets and the targets of >30% of all drugs. Because they are membrane proteins with flexible and dynamic structures, biochemical and crystallography experiments are difficult. Only ~50 GPCRs out of > 800 in the human genome have been crystallized despite their great pharmacological importance. Recent advances in homology modeling have allowed more accurate ligand docking (1), but GPCR modeling remains challenging due to conformational flexibility and abundance of flexible loops.

Various computational approaches have been used to predict ligand binding sites in G-protein coupled receptors. Traditional docking methods compute the lowest energy pose of a ligand fit to a receptor surface. Such methods are highly dependent on the form of the energy scoring function and accuracy of the receptor model structure (2–4). These methods have been used to identify ligand binding sites and build pharmacophores for G-protein coupled receptors (GPCRs) (5–7), but the lack of diverse GPCR crystal structures presents serious challenges to using docking methods for identification of ligand binding sites. Moreover, homology models usually cannot be used to identify ligand binding sites or for docking without extensive optimization (4, 8). An underappreciated feature that can be used to predict ligand binding sites is surface or sequence conservation. Binding sites for particular ligands are often conserved, although systematic sequence variation can encode ligand specificity (9–11). The massive abundance of genomic data for GPCRs can provide strong constraints for possible ligand binding sites even without chemical or structural information (12–14).

There has been less research on methods that combine information from chemical interactions, geometric surface analysis, and bioinformatics. Hybrid strategies, such as Concavity (15), have demonstrated superior performance in predicting ligand binding sites compared to single-mode approaches. Concavity scores binding sites by evolutionary sequence conservation, as quantified by the Jensen-Shannon divergence (11), and employs geometric criteria of size and shape. Here, we describe and apply a new hybrid strategy, ConDock, that combines information from surface conservation with intermolecular interactions from docking calculations. We compare our results from those previously published using purely docking-based and other hybrid methods (16, 17). We demonstrate the effectiveness of ConDock for identifying ligand binding sites for two GPCRs with known crystal structures, the β2 adrenergic and A2A adenosine receptors.

We also apply ConDock to predict the binding sites of four ligands to the less characterized G-protein coupled estrogen receptor (GPER, formerly known as GPR30), a membrane-bound estrogen receptor. GPER is proposed to mediate rapid estrogen-associated effects, cAMP regeneration, and nerve growth factor expression (18–22). GPER is known to bind estradiol and the estrogen receptor inhibitors, tamoxifen and fulvestrant, that are used to treat breast cancer (Fig. S1). Recently, GPER-specific ligands G1 and G15 were discovered (23, 24). G1 and G15 are structurally similar, differing by only an acetyl group. G1 is an agonist, whereas G15 is an antagonist. No crystal structure of GPER is available, and details of ligand binding are unknown. We discuss how the ConDock-predicted binding sites provide a basis for G1 and G15 binding specificity.

## Results

The A2A adenosine and β2 adrenergic receptors are the most heavily studied GPCRs by crystallography. We used them as standards to validate the effectiveness of ConDock for predicting ligand binding sites. For both receptors, we performed cross-docking of an agonist and inverse agonist against a crystal structure of the receptor bound to a different agonist or inverse agonist: ligands were cross-docked rather than self-docked into its own crystal structure. Docking was performed with SwissDock which has demonstrated high accuracy in docking ligands into receptors without prior knowledge of the binding site and also includes a user-friendly web interface (25). SwissDock docking results were then ranked by the ConDock scoring function (Table 1). Residues within 3.5 Å of the highest scoring predicted ligand sites were compared with the poses in the crystal structures. In addition, we determined the distances between the centers of mass for the poses in the crystal structure and those scored highest by ConDock. As negative controls, we docked morphine to the active conformations of the A2A adenosine and β2 adrenergic receptors.

**Table 1.** ConDock predicted ligand binding sites for A2A adenosine and β2 adrenergic receptors using crystal structures.

| GPCR | Ligand | Distance to crystal structure pose (Å) | ConSurf conservation score | SwissDock energy Score | ConDock score |
| --- | --- | --- | --- | --- | --- |
| A2A adenosine | adenosine (agonist) | 1.8 | 0.86 | -1702.1 | -1458.9 |
| A2A adenosine | ZM241385 (inverse agonist) | 1.0 | 0.86 | -1534.7 | -1319.8 |
| β2 adrenergic | epinephrine (agonist) | 0.4 | 0.87 | -1112.0 | -970.7 |
| β2 adrenergic | carazolol (inverse agonist) | 1.8 | 0.78 | -1534.7 | -1319.8 |

The highest ranked pose for adenosine within the A2A adenosine receptor was within 1.8 Å of the ligand position in the crystal structure. (Fig. 1A). The ConDock-predicted binding site had a ConSurf conservation score of 0.86 and is essentially the same as the experimental binding. The highest ranked site for ZM241385 within the A2A adenosine receptor was within 1. 0 Å of the ligand’s position in the crystal structure. In this top pose, ZM241385 is found within the same binding site as that observed in the crystal structure (Fig. 1B), with a ConSurf conservation score of 0.86.

**Figure 1.**
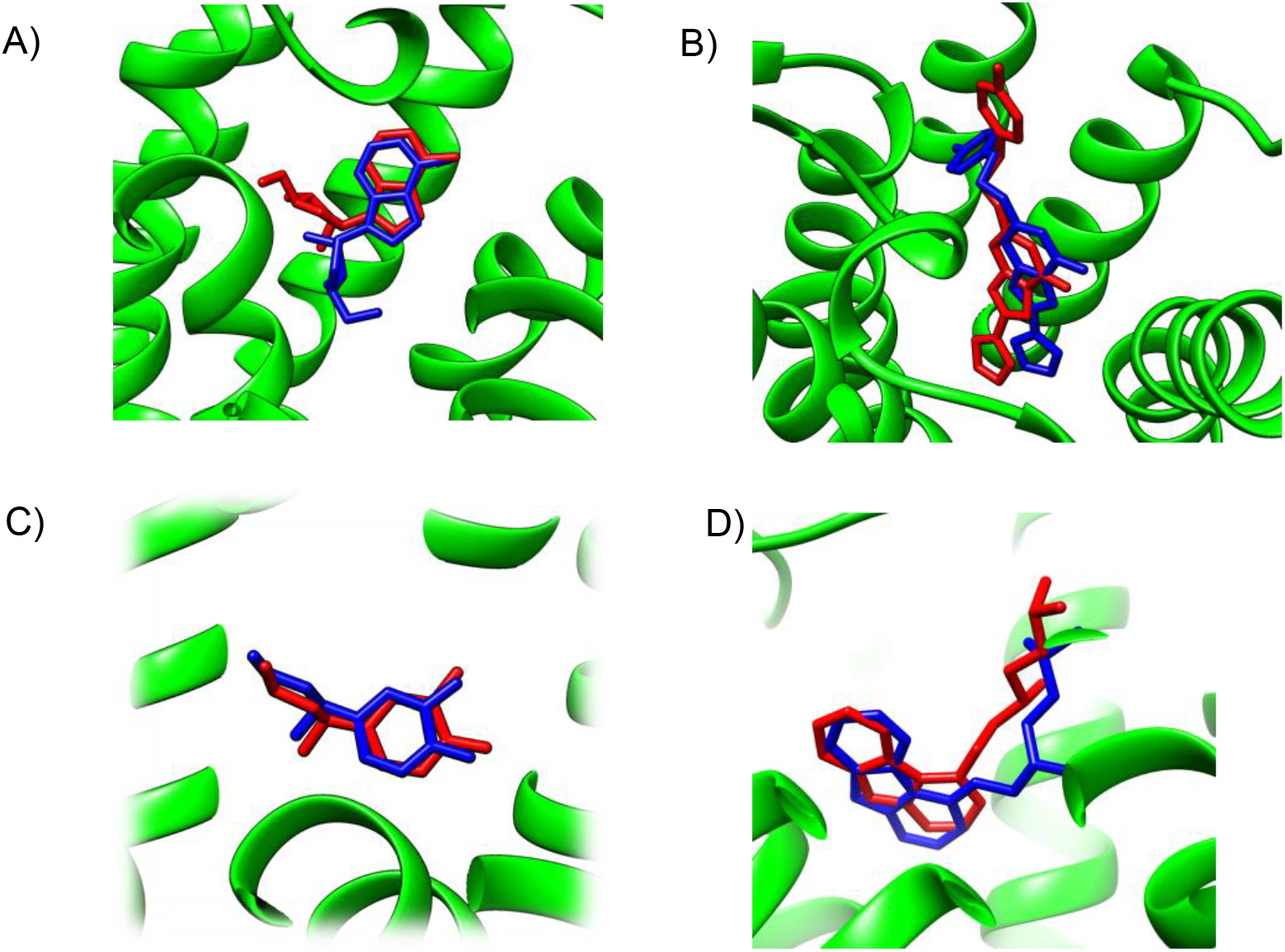
Predicted and experimental ligand binding sites in A2A adenosine and β2 adrenergic receptors. Superposition of crystal structure with ligand bound (red) with ConDock predicted pose (blue). A) Adenosine with A2A receptor. B) ZM241385 with A2A receptor. C) Epinephrine with β2 adrenergic receptor. D) Carazolol with β2 adrenergic receptor.

The highest ranked pose for epinephrine within the β2 adrenergic receptor was within 0.4 Å of the ligand position within the crystal structure (4LDO). This binding site for epinephrine was again essentially the same as the observed binding pocket (Fig. 1C). The highest ranked pose for carazolol within the β2 adrenergic receptor was within 1.8 Å of the ligand’s position within the crystal structure (2RH1). This binding site for carazolol was essentially the same as that in the crystal structure (Fig. 1D). This pose possessed a ConSurf conservation score of 0.78. The extremely accurate placement of both agonists and antagonists demonstrates ConDock’s effectiveness when a GPCR crystal structure is available. For our negative control case, SwissDock failed to dock morphine within the ligand binding cavity of the receptors at all (Fig. S1). This demonstrates the utility of including docking in a hybrid strategy to eliminate false positives for ligand binding sites.

Unfortunately, crystal structures are not available for many GPCRs. The most valuable use of ConDock is predicting drug binding sites in homology models. By using surface conservation information, ConDock is potentially less sensitive to homology model inaccuracies than other ligand binding site prediction methods that are based purely on geometric methods. To demonstrate the ability of ConDock to work with homology models, we created models of four GPCRs, the β2 adrenergic, A2A adenosine, 5HT2B serotonin, and mu opioid receptors, that did not use the known crystal structures as templates. We used I-TASSER (26) for modeling. I-TASSER created fairly accurate models of all four receptors, with rmsds between the models and crystal structures ranging from a best of 0.85 Å for the β2 adrenergic receptor (pdb 2rh1) to a respectable 2.1 Å for the A2A adenosine receptor (pdb 5k2a). We used ConDock to predict the binding sites of the β2 adrenergic receptor with carazalol, A2A adenosine receptor with ZM241385, 5HT2B serotonin receptor with methysergide, and mu opioid receptor with BU72.

Not surprisingly, ConDock performed best with the highly accurate β2 adrenergic homology model, with only 1.8 Å between the predicted and crystal structure ligand poses centers of mass (Table 2, Fig. 2). Performance decreases for the other three receptors with less reliable homology models. The A2A adenosine and mu opioid receptors (pdb 5c1m) have distances of 3-4 Å between predicted and crystal structure ligand poses. In this distance range, most of the residues are the same between the predicted and crystal structure binding sites, supporting a successful prediction. ConDock performs less well with the 5HT2B serotonin receptor (pdb 6drz) for which the distance between the predicted and actual ligand binding sites was 7.3 Å. In the 5HT2B receptor structure, the ligand, methysergide, binds very deep in the receptor. Inaccurate modeling of the receptor makes it difficult or impossible to dock the ligand into the deep, restricted binding site.

**Figure 2.**
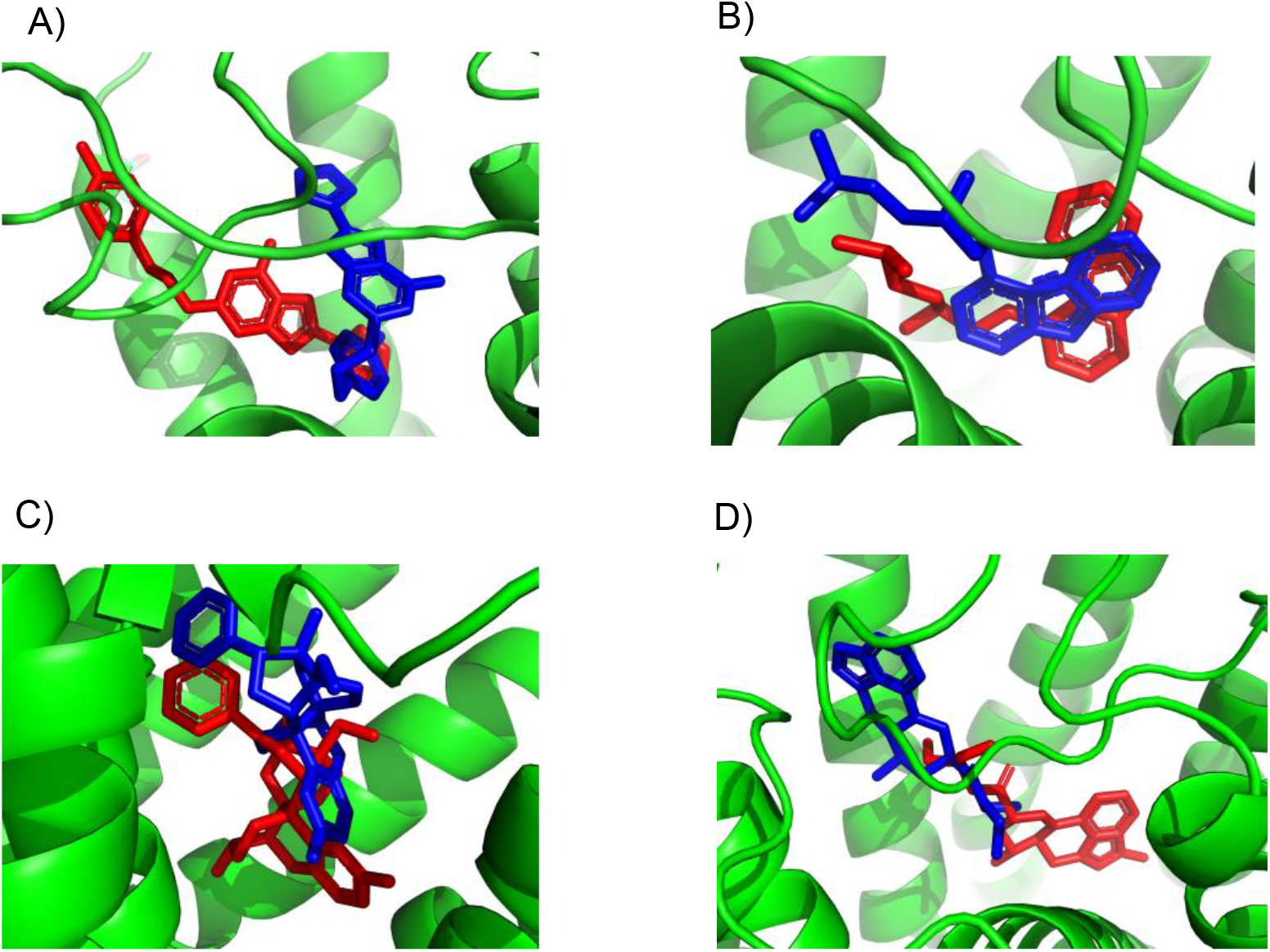
Predicted and experimental ligand binding sites for homology models of four GPCRs. Superposition of crystal structure with ligand bound (red) with ConDock predicted pose (blue). A) ZM241385 with A2A adenosine receptor. B) Carazolol with β2 adrenergic receptor. C) BU72 with mu opioid receptor. D) Methysergide with 5HT2B serotonin receptor.

**Table 2.**
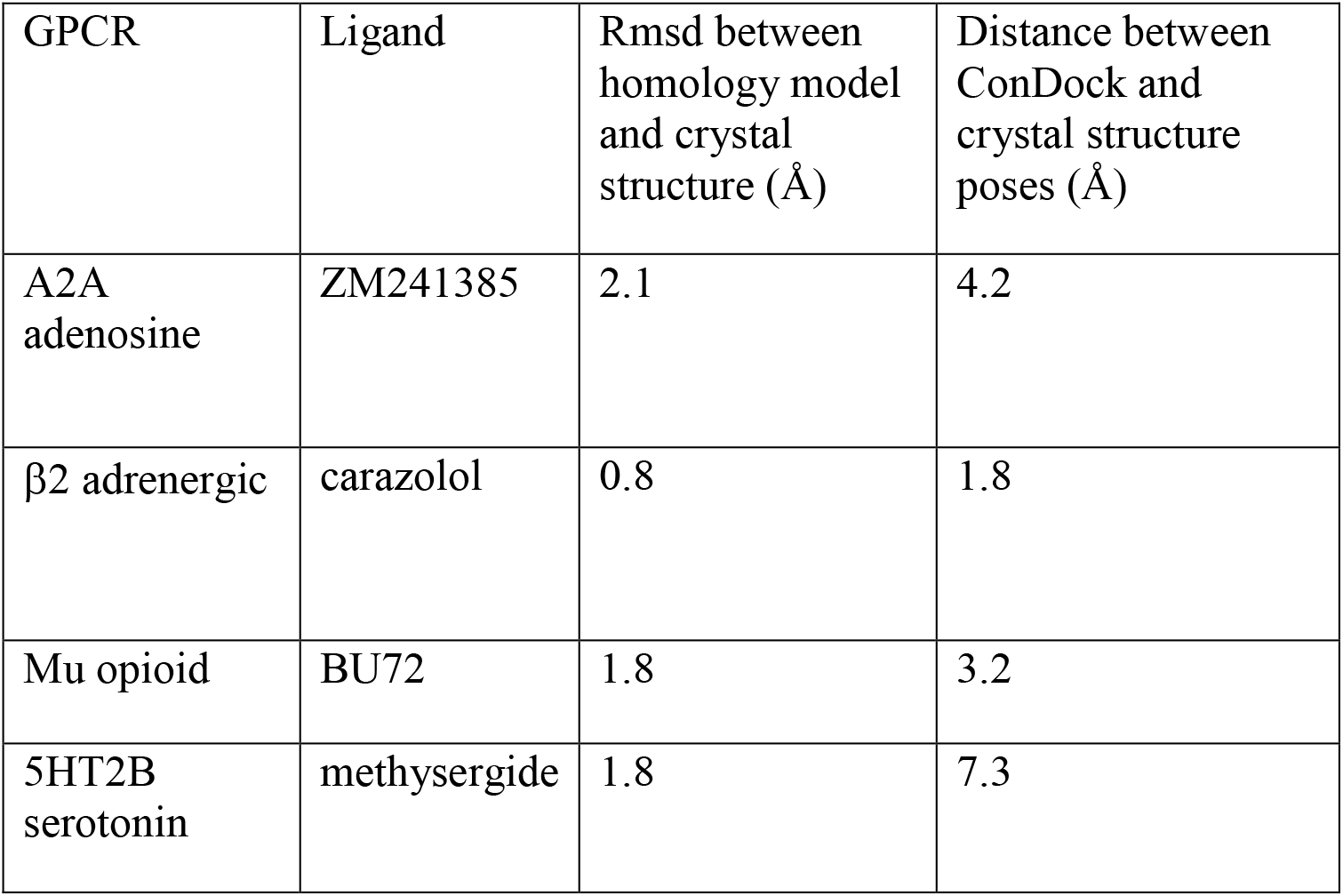
ConDock predicted ligand binding sites for four GPCRs using homology models.

| GPCR | Ligand | Rmsd between homology model and crystal structure (Å) | Distance between ConDock and crystal structure poses (Å) |
| --- | --- | --- | --- |
| A2A adenosine | ZM241385 | 2.1 | 4.2 |
| $\beta 2$ adrenergic | carazolol | 0.8 | 1.8 |
| Mu opioid | BU72 | 1.8 | 3.2 |
| 5HT2B serotonin | methysergide | 1.8 | 7.3 |

After demonstrating the accuracy of ConDock for high quality homology models, we applied ConDock to predict the binding sites in a GPCR, GPER (G-protein coupled estrogen receptor), which has not yet been crystallized. To predict the potential ligand binding sites in GPER, we first created a homology model using GPCR-I-TASSER (27). GPCR-I-TASSER has been shown to be the most accurate GPCR homology modeling software package. GPCR-I-TASSER identified the closest matching crystal structure to GPER to be the CCR5 chemokine receptor (PDB 4mbs) with 23% sequence identity. GPCR-I-TASSER used this crystal structure along with 9 other GPCR crystal structures as templates for homology modeling. The GPER homology model differs from chain A of the crystal structure of CCR5 chemokine receptor with RMSD of 0.96 Å across Cα atoms (Fig. S3) and has an excellent Ramachandran plot (Fig. S4). The primary differences are in the extracellular loop between helices 4 and 5 and the intracellular loops between helices 5 and 6, and after helix 7. These two intracellular loops are predicted by ERRAT (28) to be the least reliable based on the likelihood of atom pair type interactions from high-resolution crystal structures (Fig. S5). As the sites with the greatest predicted errors are on the intracellular face of GPER, far from the ligand binding site, they are less likely to affect our ligand prediction study, giving us greater confidence in our homology model.

Using the SwissDock server (25), we docked structures of the four ligands E2, G1, G15, and tamoxifen (Fig. S1) to the homology model of GPER. Most of the docked sites from SwissDock were not located on the extracellular face of GPER and thus were considered nonviable (Fig. S6). The shortcomings of a purely physics-based scoring function such as that used by SwissDock in predicting ligand binding is not surprising given the lack of an experimental crystal structure and well-known limitations of current computational methodology (29–32).

We then ranked all ligand binding sites generated by SwissDock using the combined ConDock score. The ConDock score is simply the product of the ConSurf (33, 34) binding surface sequence conservation score and the SwissDock FullFitness energy score (35). A highly negative ConDock score is associated with a more probable ligand binding site. For all four ligands, the ConDock score identified one or two ligand binding sites and poses that clearly outscored other candidates (Table 3). ConDock identified the same approximate binding site for all four ligands, although this was not an explicit criterion in the calculations (Fig. 3). The average ConSurf conservation score across the four ligand binding sites is 0.82 (1.0 represents complete conservation), indicating that the site is highly but not completely conserved. The binding site is located deep in the receptor cleft, although depth was not a criterion in the prediction calculation. Given the lack of additional experimental evidence for the location of the ligand binding site, the proposed ConDock sites are physically reasonable.

**Table 3.**
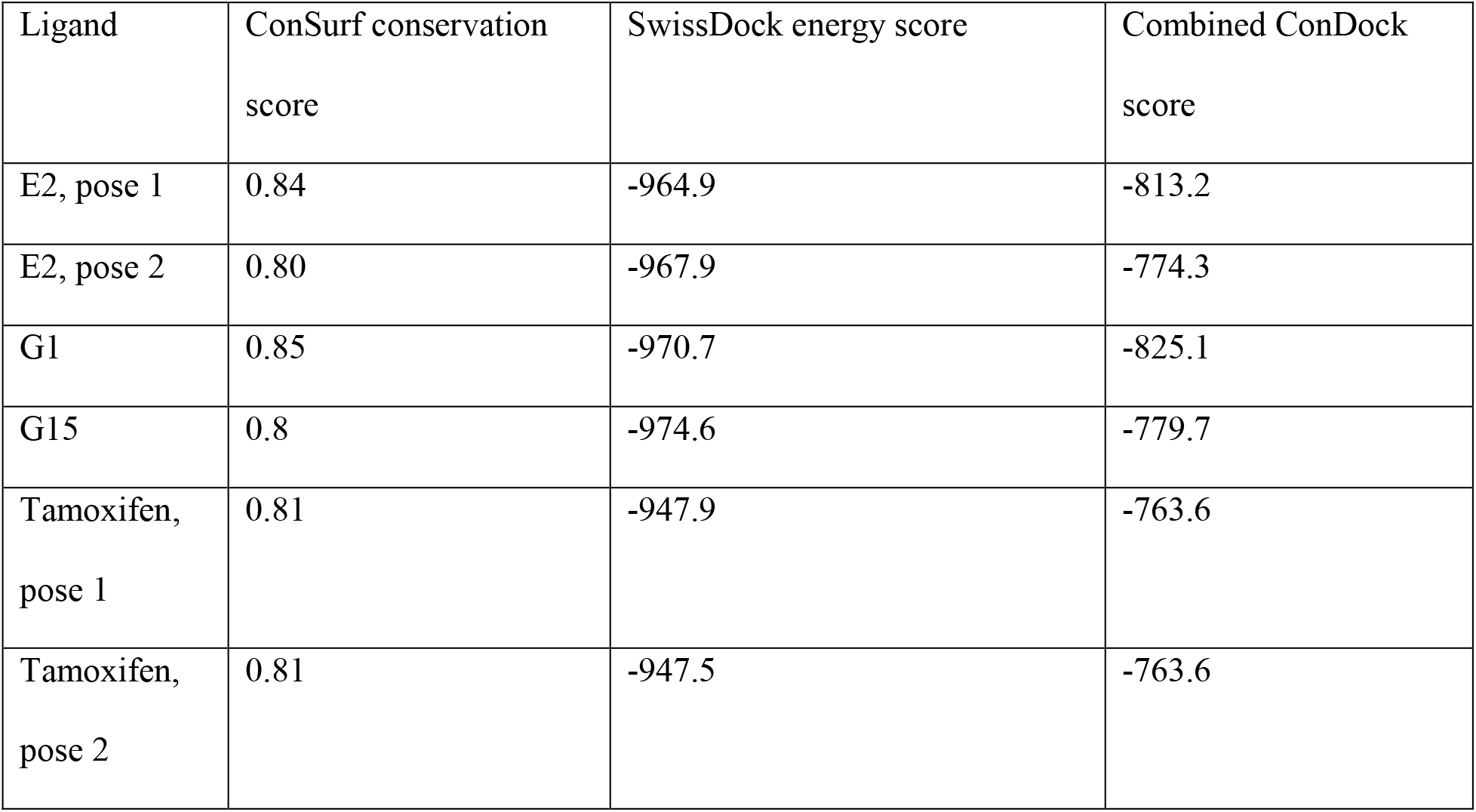
Scores for predicted binding sites and poses for GPER ligands.

**Figure 3.**
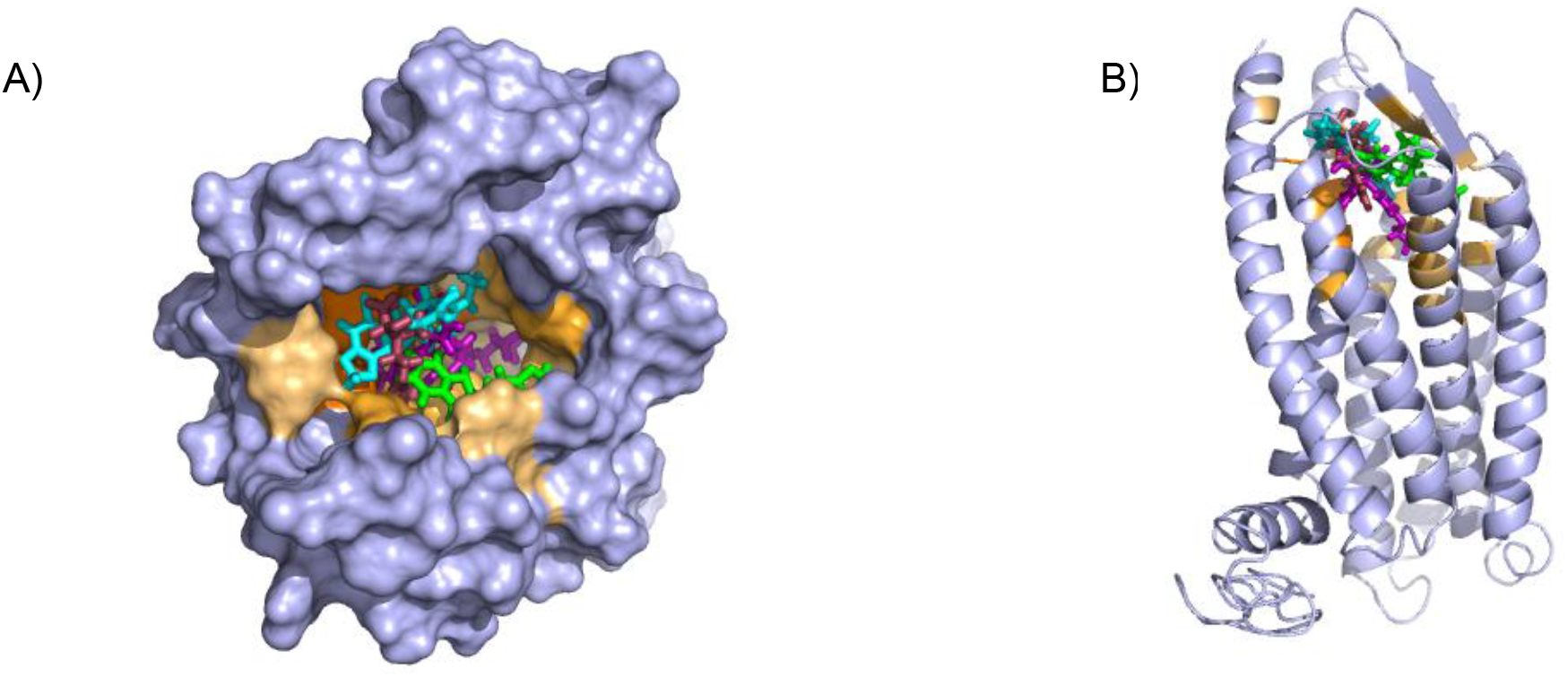
Predicted ligand binding pockets in GPER. A) Extracellular perspective of GPER showing amino acids predicted to contact ligands E2 (maroon), G1 (cyan), G15 (green), and tamoxifen (violet). Residues colored orange are predicted to contact one or more ligands with darker hue indicating interaction with multiple ligands. B) Predicted binding pocket viewed from a 90° rotation.

We found two promising binding sites for E2 in GPER. The two sites are 4.4 Å apart, located deep in the receptor cleft (Fig. 3). E2 is oriented perpendicular to the lipid membrane and rotated about 180° between the two poses. The conservation scores for these two poses are 0.84 and 0.80. The energy scores of the two poses are similar. The amino acids contacting E2 in pose 1 are conserved in GPERs from six species, and only one residue contacting pose 2, H282, varies across species. In the top ranked pose, there is a hydrogen bond between the inward pointing D-ring hydroxyl group of E2 and the carboxyl terminal on E115. Hydrophobic interactions are present between E2 and non-polar residues L119, Y123, P303, and F314. In the second ranked pose, the inward pointing A-ring hydroxyl group of E2 makes a hydrogen bond with N310. This pose is in a less hydrophobic environment, contacting primarily H282 and P303.

ConDock predicts that G1 and G15 bind in adjacent but distinct binding sites separated by 2.3 Å despite the chemical similarity of the two ligands. The top predicted binding site for G1 is found within the pocket bound by Y55, L119, F206, Q215, I279, P303, H307, and N310 (Fig. 4). This orientation had the highest conservation score of all predicted binding sites at 0.85. In this pose, N310 makes a long hydrogen bond with the acetyl oxygen of G1. The predicted binding site for G15 is found within the pocket bound by L119, Y123, M133, S134, L137, Q138, P192, V196, F206, C207, F208, A209, V214, E218, H307, and N310. This pose had a conservation score of 0.8. Hydrogen bonding is not observed between GPER and G15. Hydrophobic interactions are observed with L119, Y123, F206, and V214.

**Figure 4.**
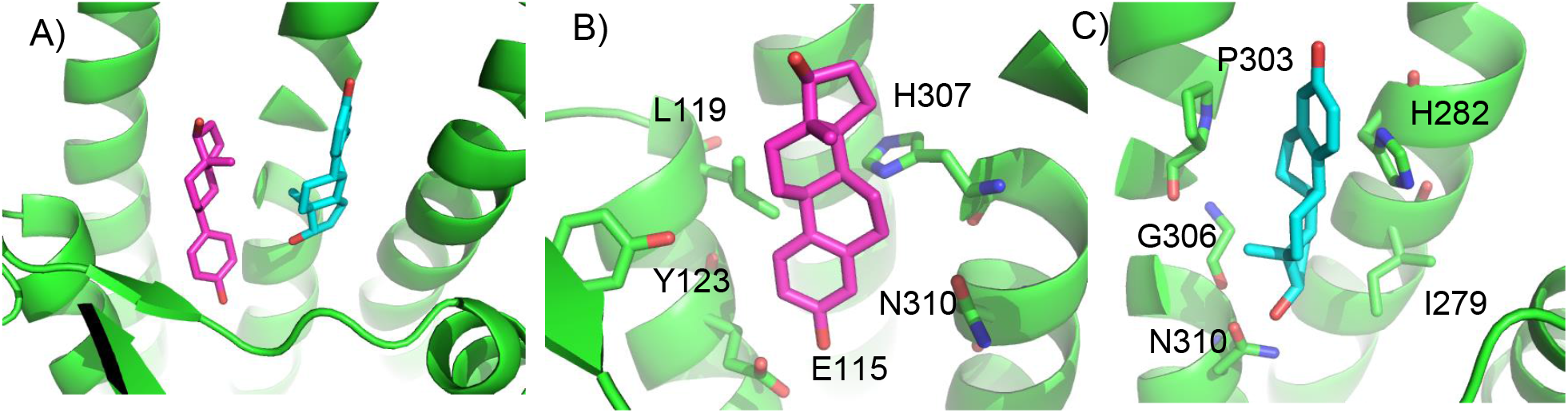
Predicted E2 binding sites in GPER. A) The two highest scoring docking poses for E2. B) Receptor-ligand interactions for E2 pose 1. C) Receptor-ligand interactions for E2 pose 2.

ConDock predicted two equally high-scoring, overlapping poses for tamoxifen, near E115, L119, Y123, L137, Q138, M141, Y142, Q215, E218, W272, E275, I279, P303, G306, H307, and N310 (Fig. 5). The conservation score of this orientation is 0.81. Hydrophobic interactions are observed between tamoxifen and non-polar residues L119, Y123, Y142, P303, and F314. Notably, the amine group of tamoxifen is neutralized by E218 and E275.

**Figure 5.**
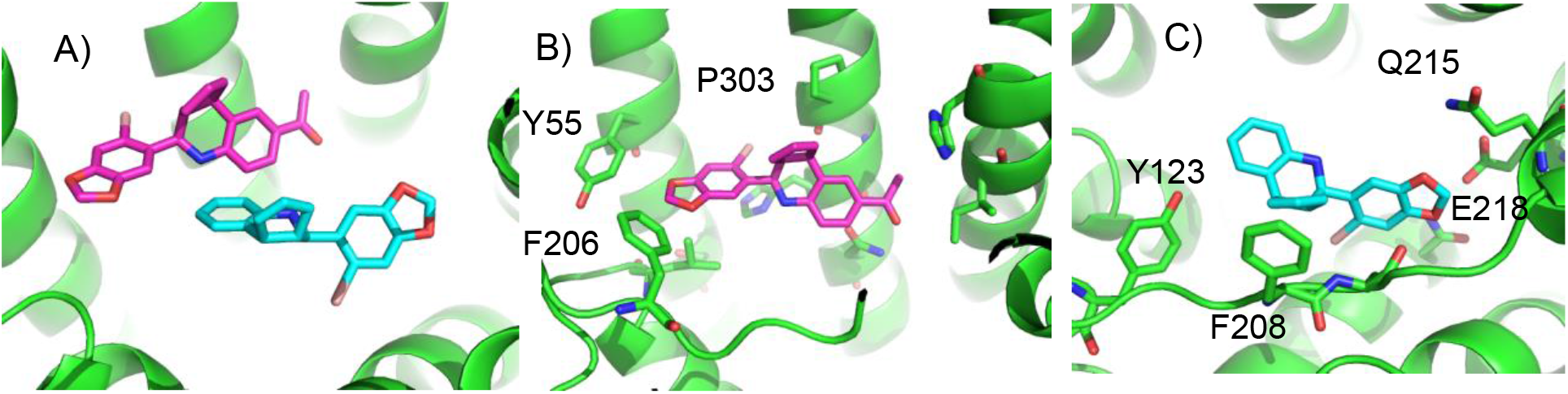
Predicted G1 and G15 binding sites in GPER. A) The highest scoring docking poses for G1 (maroon) and G15 (cyan). B) Receptor-ligand interactions for G1. C) Receptor-ligand interactions for G15.

In comparison with the ligand binding sites predicted by traditional methods based on surface geometry and conservation (Fig. S7), the sites predicted by ConDock are more detailed and of higher resolution due to the information from chemical interactions from ligand docking. Moreover, prediction methods based on surface geometry and conservation cannot differentiate between binding sites for different ligands. We compared the GPER ligand binding sites predicted by ConDock to those predicted by three other software packages representing different approaches: CASTp (36), which analyzes surface geometry, SiteHound (37), which maps surfaces with a chemical probe, and Concavity (15), which analyzes surface geometry and conservation (Fig. 6). All three methods could identify a ligand binding site roughly matching that from ConDock. The pocket predicted by ConDock is deeper than the other pockets, which while intuitively attractive, is not necessarily correct. SiteHound performed particularly poorly, with the top scoring site located on the GPER intracellular face. The site identified by SiteHound closest to the ConDock site was scored third and is a shallow binding pocket near H52-G58, E275-H282, and R299-H307 (Fig. 6C). In contrast, the Concavity site was smaller and shallower than the ConDock site (Fig. 6D). Surprisingly, the site predicted by the simpler CASTp method best matched the ConDock site but is also smaller and shallower (Fig. 6B). For proteins such as GPCRs with large, concave binding pockets, geometry-based prediction methods such as Concavity and CASTp can easily identify the general location of the binding site. However, such methods may have more difficulty recovering the specific, ligand-specific binding site. It is also surprising that ConDock more closely matched the results of the geometry-based methods given that ConDock does not take surface geometry into account. Without experimental structural data, it is not possible to conclude which of the predicted binding sites is correct at this time.

**Figure 6.**
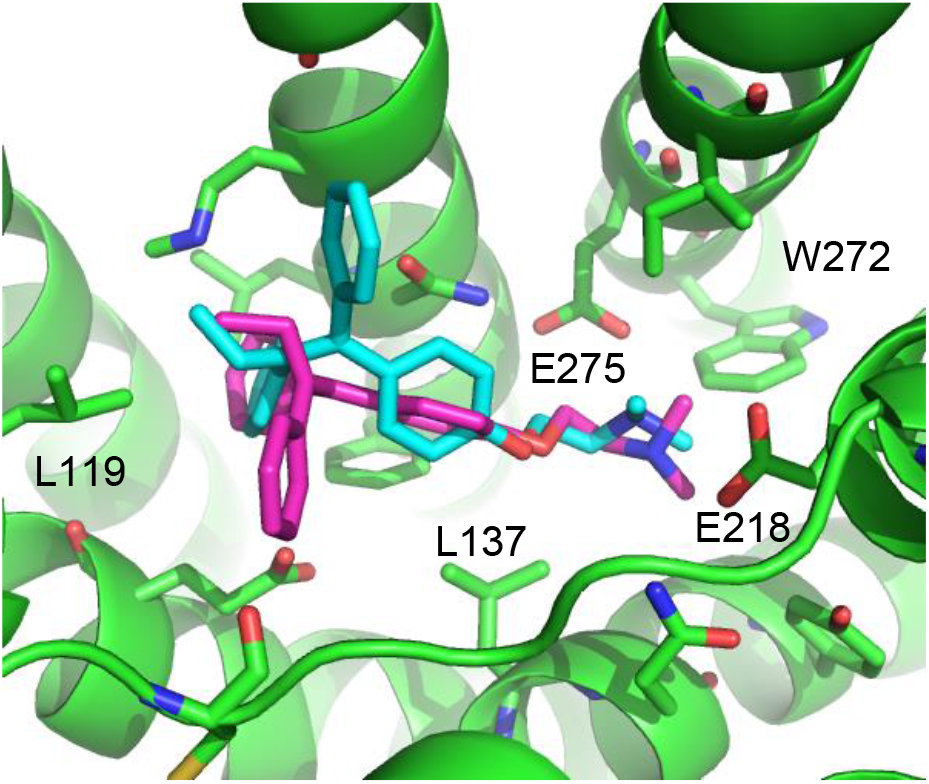
Predicted tamoxifen binding sites in GPER. A) The highest scoring docking poses for tamoxifen, pose 1 (maroon) and pose 2 (cyan).

## Discussion

The ConDock scoring method, incorporating information from both surface conservation and docking binding energy, demonstrated high accuracy in predicting ligand binding sites from the crystal structures of the A2A adenosine and β2 adrenergic receptors. ConDock also successfully predicted the ligand binding sites from high-quality homology models. ConDock was also used to predict viable ligand binding sites for four different GPER ligands. In contrast to more typical geometry-based ligand binding site prediction methods, ConDock scoring takes advantage of chemistry-specific information about the ligand-receptor interface. The poor performance of SiteHound in predicting ligand binding sites on GPER suggests that a method based only on chemical interactions or docking is highly susceptible to error, most likely due to the inadequate accuracy of homology models. Surface conservation data not only provides orthogonal knowledge but also dampens the influence from the shortcomings of current computational methods in homology modeling, docking, and predicting binding affinity. How best to mathematically combine these multiple data sources has been debated (11, 15), but we demonstrate here that a simple product scoring function is effective. The four GPER ligands analyzed differ greatly in chemical structure, but the ConDock scoring method predicted that all four bind to the same approximate region, deep in the extracellular cleft of the receptor. Undoubtedly, further refinement of a hybrid scoring function will lead to improved predictions.

Recent GPER modeling studies using molecular dynamics simulations and docking identified different potential binding sites for E2, G1, and G15 near F206 and F208; the interaction with this region was described as driven primarily by π-π stacking interactions (16, 17). Figure 6 compares the ConDock binding site against that predicted in the molecular dynamics simulation and docking study. The ConDock binding site is located deeper in the extracellular cleft; the other proposed site mostly involved surface-exposed loops. Mendez-Luna *et al.* proposed that Q53, Q54, G58, C205, and H282 all interact with G1 and G15; however, none of these residues are conserved across the six species we analyzed. Experimental data is not currently available to support one model over the other.

In summary, the simple ConDock hybrid scoring model predicts physically plausible ligand binding sites by combining information from ligand docking and surface conservation. Using multiple orthogonal sources of information avoids errors introduced by modeling, especially in a case where a crystal structure of the receptor is unavailable. Given a high quality homology model, ConDock can accurately predict ligand binding sites. Using this hybrid method, we identified a site in the extracellular cleft of GPER that has the potential to bind four known GPER ligands. Further optimization of hybrid scoring functions should yield significantly improved predictions.

## Methods

### Protein surface conservation

GPCR protein sequences were acquired from the SwissProt database (38). For the A2A adenosine receptor, the protein sequences aligned were from *Homo sapiens, Canis familiaris, Xenopus tropicalis, Myotis davidii, Loxodonta africana, Gallus gallus, Anolis caronlinesis, Oncorhynchus mykiss, Ailuropoda melanoleuca*, and *Alligator mississippiensis*. For the β2 adrenergic receptor, the protein sequences aligned were from *Homo sapiens, Oncorhynchus mykiss*, *Myotis brandtii*, *Callorhinchus milii*, *Ophiophagus hannah*, *Canis familiaris*, *Loxodonta africana, Ailuropoda melanoleuca, Ficedula albicollis*, and *Xenopus laevis.* GPER protein sequences aligned were from diverse species: *Homo sapiens*, *Rattus norvegicus*, *Mus musculus*, *Macaca mulatta, Danio rerio*, and *Micropogonias undulatus.* Sequences were chosen to represent a diverse range of animal species. Multiple sequence alignment files were submitted to ConSurf (33, 34). ConSurf assesses conservation using Bayesian reconstruction of a phylogenetic tree. Each sequence position is scored from 0-9, where 9 indicates that the amino acid was retained in all the organisms (Fig. S7). Values from ConSurf were mapped onto the receptor surface with Chimera (39).

### Homology modeling and docking

The crystal structures for the A2A adenosine receptor and the β2 adrenergic receptor were acquired from the RCSB protein data bank: β2 adrenergic receptor bound to epinephrine (PDB 4ldo), β2 adrenergic receptor bound to carazolol (PDB 2rh1), A2A adenosine receptor bound to adenosine (PDB 2ydo), and A2A adenosine receptor bound to ZM241385 (PDB 5k2a). The crystal structures of the mu opioid receptor and 5HT2B receptor were taken from PDB 5c1m and 6drz. Structures were prepped for docking with Chimera by removing extraneous chains and bound ligands with the DockPrep protocol. Ligands were docked into receptors with SwissDock (25).

The crystal structure of GPER has not yet been determined. We created a homology model using GPCR I-TASSER (Iterative Threading Assembly Refinement), the most accurate homology modeling software customized for GPCRs (27). GPCR I-TASSER modeled the GPER structure using templates from the ten closest related GPCR crystal structures (PDB 4mbs, 2ks9, 1kpn, 1u19, 2ziy, 1kp1, 3odu, 4ea3, 4iaq, 2y00). The homology model was validated with ERRAT (28). Coordinates for E2, G1, G15, and tamoxifen were downloaded from the ZINC ligand database (40) and submitted to SwissDock (25) for docking. SwissDock is a web interface to the EADock DSS (35) engine, which performs blind, global (does not require targeting of a particular surface) docking using the physics-based CHARMM22 force field (41). The “FullFitness Score” calculated by SwissDock using clustering and the FACTS implicit solvent model (42) was used as the “Energy Score” for our calculations.

### Combined analysis

SwissDock poses were manually screened for those binding sites located on or near the extracellular side of the protein. Ligand binding surfaces included residues with atoms within 3.5 Å from the docked ligand. The average conservation score of the amino acids that were highlighted served as the “Conservation Score” of that specific orientation (Fig. 8). The combined ConDock score is defined as the product of the Conservation and Energy Scores. As the Energy Score is a modified free energy function, a highly negative ConDock score is associated with a more probable ligand binding site. Binding sites predicted by ConDock results were compared with those predicted by CASTp (36), SiteHound (37), and Concavity (15). For CASTp, SiteHound, and Concavity, ligand binding pockets were defined as residues within 4 Å of the selected probe/cluster.

**Figure 7.**
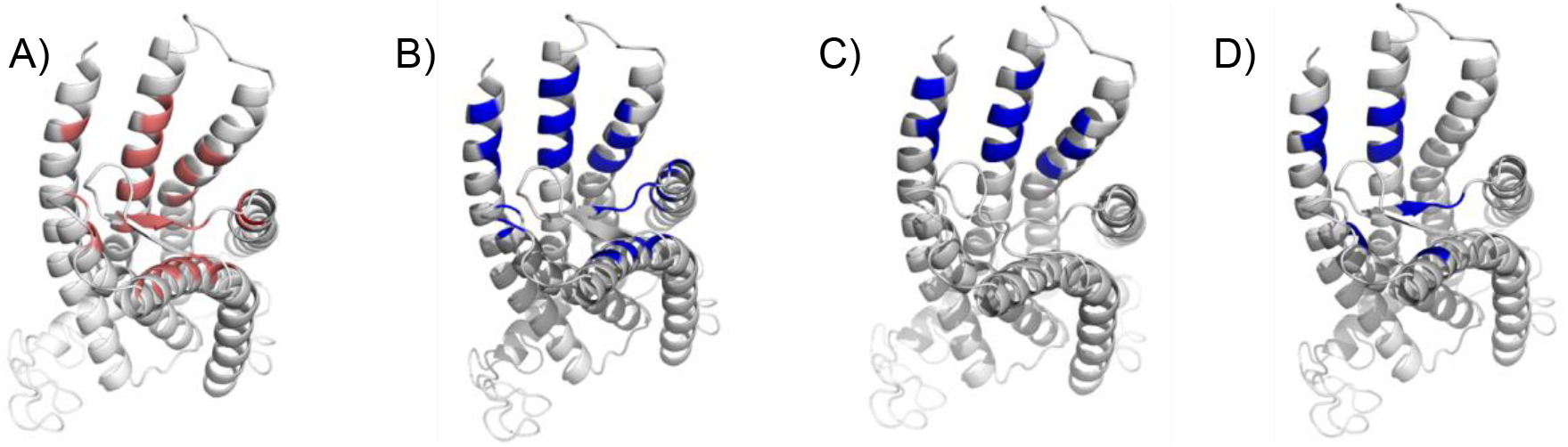
Predicted ligand binding sites by ConDock, CASTp, SiteHound, Concavity. Ligand binding sites are colored, predicted by A) ConDock, B) CASTp, C) SiteHound, D) Concavity.

**Figure 8.**
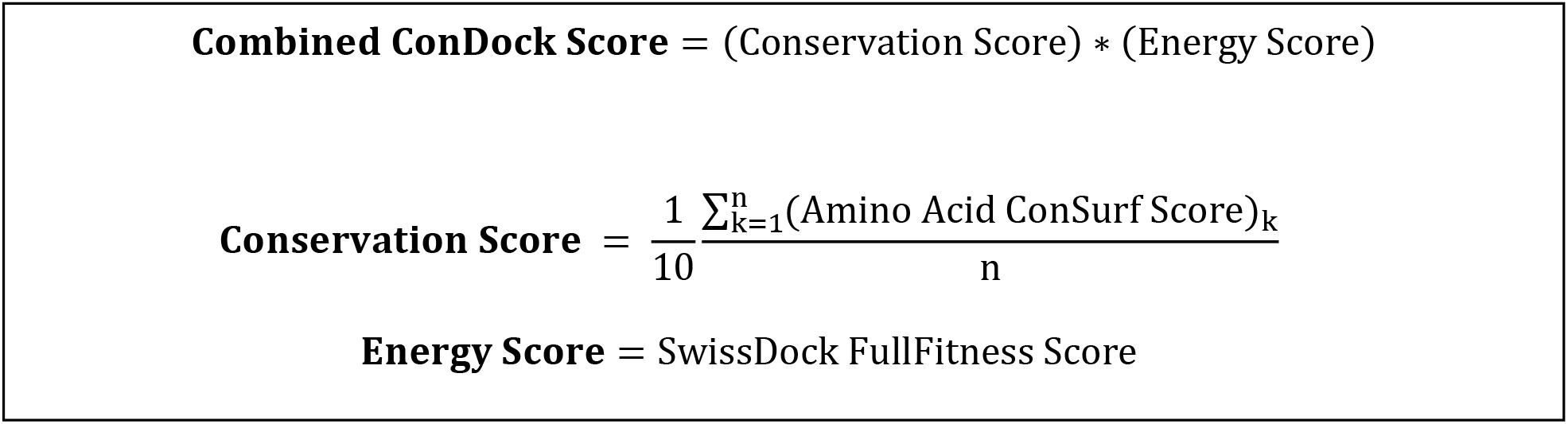
Calculation of combined ConDock scores for ligand binding sites. The Conservation Score is calculated over the *n* residues in a binding site, indexed by *k*.

### Crystal structure benchmarks

Crystal structures of receptors were screened for residues within 3.5 Å of their respective ligands. These residues served as a benchmark of comparison for the predicted sites contrived by the ConDock scoring function.

## Acknowledgments

This work was funded by the Victoria S. and Bradley L. Geist Foundation (H.L.N.), NSF CAREER Award 1350555 (H.L.N.), and the Undergraduate Research Opportunities Program at the University of Hawaii at Manoa (A.R.V., S.M.).

## Author contributions

A.R.V., S.M., and H.L.N. performed the analysis and calculations. A.R.V., S.M., and H.L.N. wrote the manuscript. H.L.N. supervised the project.

## Competing financial interests

The authors have no competing financial interests.

## Notes

#### Summary of Updates

Added section on analyzing homology models.

## References

1. Feng X, Ambia J, Chen K-YM, Young M, Barth P (2017) Computational design of ligand-binding membrane receptors with high selectivity. Nat Chem Biol 13(7):715–723.

2. Shoichet BK, Kobilka BK (2012) Structure-based drug screening for G-protein-coupled receptors. Trends Pharmacol Sci 33(5):268–272.

3. Katritch V, Abagyan R (2011) GPCR agonist binding revealed by modeling and crystallography. Trends Pharmacol Sci 32(11):637–643.

4. Katritch V, Rueda M, Lam PC-H, Yeager M, Abagyan R (2010) GPCR 3D homology models for ligand screening: lessons learned from blind predictions of adenosine A2a receptor complex. Proteins 78(1):197–211.

5. Sanders MPA, et al. (2011) Snooker: A Structure-Based Pharmacophore Generation Tool Applied to Class A GPCRs. J Chem Inf Model 51(9):2277–2292.

6. Kratochwil NA, et al. (2011) G protein-coupled receptor transmembrane binding pockets and their applications in GPCR research and drug discovery: a survey. Curr Top Med Chem 11(15):1902–1924.

7. Tang H, Wang XS, Hsieh J-H, Tropsha A (2012) Do crystal structures obviate the need for theoretical models of GPCRs for structure-based virtual screening? Proteins Struct Funct Bioinforma 80(6): 1503–1521.

8. Lai JK, Ambia J, Wang Y, Barth P (2017) Enhancing Structure Prediction and Design of Soluble and Membrane Proteins with Explicit Solvent-Protein Interactions. Structure 25(11):1758–1770.e8.

9. Wass MN, Sternberg MJE (2009) Prediction of ligand binding sites using homologous structures and conservation at CASP8. Proteins Struct Funct Bioinforma 77(S9): 147–151.

10. Kalinina OV, Gelfand MS, Russell RB (2009) Combining specificity determining and conserved residues improves functional site prediction. BMC Bioinformatics 10:174.

11. Capra JA, Singh M (2007) Predicting functionally important residues from sequence conservation. Bioinformatics 23(15):1875–1882.

12. Madabushi S, et al. (2004) Evolutionary Trace of G Protein-coupled Receptors Reveals Clusters of Residues That Determine Global and Class-specific Functions. J Biol Chem 279(9):8126–8132.

13. Levit A, Barak D, Behrens M, Meyerhof W, Niv MY (2012) Homology model-assisted elucidation of binding sites in GPCRs. Methods Mol Biol Clifton NJ 914:179–205.

14. Sanders (2011) ss-TEA: Entropy based identification of receptor specific ligand binding residues from a multiple sequence alignment of class A GPCRs. BMC Bioinformatics 12(Suppl 6):332–343.

15. Capra JA, Laskowski RA, Thornton JM, Singh M, Funkhouser TA (2009) Predicting Protein Ligand Binding Sites by Combining Evolutionary Sequence Conservation and 3D Structure. PLoS Comput Biol 5(12):e1000585.

16. Méndez-Luna D, et al. (2015) Deciphering the GPER/GPR30-agonist and antagonists interactions using molecular modeling studies, molecular dynamics, and docking simulations. J Biomol Struct Dyn 33(10):2161–2172.

17. Arnatt CK, Zhang Y (2013) G Protein-Coupled Estrogen Receptor (GPER) Agonist Dual Binding Mode Analyses Toward Understanding of Its Activation Mechanism: A Comparative Homology Modeling Approach. Mol Inform 32(7):647–658.

18. Carmeci C, Thompson DA, Ring HZ, Francke U, Weigel RJ (1997) Identification of a Gene (GPR30) with Homology to the G-Protein-Coupled Receptor Superfamily Associated with Estrogen Receptor Expression in Breast Cancer. Genomics 45(3):607–617.

19. Kvingedal AM, Smeland EB (1997) A novel putative G-protein-coupled receptor expressed in lung, heart and lymphoid tissue. FEBS Lett 407(1):59–62.

20. O’Dowd BF, et al. (1998) Discovery of Three Novel G-Protein-Coupled Receptor Genes. Genomics 47(2):310–313.

21. Filardo EJ, Quinn JA, Frackelton AR, Bland KI (2002) Estrogen Action Via the G Protein-Coupled Receptor, GPR30: Stimulation of Adenylyl Cyclase and cAMP-Mediated Attenuation of the Epidermal Growth Factor Receptor-to-MAPK Signaling Axis. Mol Endocrinol 16(1):70–84.

22. Kanda N, Watanabe S (2003) 17β-Estradiol Enhances the Production of Nerve Growth Factor in THP-1-Derived Macrophages or Peripheral Blood Monocyte-Derived Macrophages. J Invest Dermatol 121(4):771–780.

23. Bologa CG, et al. (2006) Virtual and biomolecular screening converge on a selective agonist for GPR30. Nat Chem Biol 2(4):207–212.

24. Dennis MK, et al. (2009) In vivo effects of a GPR30 antagonist. Nat Chem Biol 5(6):421–427.

25. Grosdidier A, Zoete V, Michielin O (2011) SwissDock, a protein-small molecule docking web service based on EADock DSS. Nucleic Acids Res 39(suppl):W270–W277.

26. Yang J, et al. (2015) The I-TASSER Suite: protein structure and function prediction. Nat Methods 12(1):7–8.

27. Zhang J, Yang J, Jang R, Zhang Y (2015) GPCR-I-TASSER: A Hybrid Approach to G Protein-Coupled Receptor Structure Modeling and the Application to the Human Genome. Structure 23(8):1538–1549.

28. Colovos C, Yeates TO (1993) Verification of protein structures: patterns of nonbonded atomic interactions. Protein Sci Publ Protein Soc 2(9):1511–1519.

29. Merz KM (2010) Limits of Free Energy Computation for Protein-Ligand Interactions. J Chem Theory Comput 6(5):1769–1776.

30. Wan S, Knapp B, Wright DW, Deane CM, Coveney PV (2015) Rapid, Precise, and Reproducible Prediction of Peptide–MHC Binding Affinities from Molecular Dynamics That Correlate Well with Experiment. J Chem Theory Comput 11(7):3346–3356.

31. Li Y, Hou T, Goddard III W (2010) Computational Modeling of Structure-Function of G Protein-Coupled Receptors with Applications for Drug Design. Curr Med Chem 17(12):1167–1180.

32. Smith RD, et al. (2015) CSAR Benchmark Exercise 2013: Evaluation of Results from a Combined Computational Protein Design, Docking, and Scoring/Ranking Challenge. J Chem Inf Model. doi:10.1021/acs.jcim.5b00387.

33. Armon A, Graur D, Ben-Tal N (2001) ConSurf: an algorithmic tool for the identification of functional regions in proteins by surface mapping of phylogenetic information1. J Mol Biol 307(1):447–463.

34. Ashkenazy H, Erez E, Martz E, Pupko T, Ben-Tal N (2010) ConSurf 2010: calculating evolutionary conservation in sequence and structure of proteins and nucleic acids. Nucleic Acids Res 38(Web Server issue):W529–533.

35. Grosdidier A, Zoete V, Michielin O (2011) Fast docking using the CHARMM force field with EADock DSS. J Comput Chem 32(10):2149–2159.

36. Dundas J, et al. (2006) CASTp: computed atlas of surface topography of proteins with structural and topographical mapping of functionally annotated residues. Nucleic Acids Res 34(Web Server issue):W116–118.

37. Hernandez M, Ghersi D, Sanchez R (2009) SITEHOUND-web: a server for ligand binding site identification in protein structures. Nucleic Acids Res 37(suppl 2):W413–W416.

38. Boeckmann B, et al. (2005) Protein variety and functional diversity: Swiss-Prot annotation in its biological context. C R Biol 328(10-11):882–899.

39. Pettersen EF, et al. (2004) UCSF Chimera--a visualization system for exploratory research and analysis. J Comput Chem 25(13):1605–1612.

40. Irwin JJ, Sterling T, Mysinger MM, Bolstad ES, Coleman RG (2012) ZINC: A Free Tool to Discover Chemistry for Biology. J Chem Inf Model 52(7):1757–1768.

41. Brooks BR, et al. (2009) CHARMM: the biomolecular simulation program. J Comput Chem 30(10):1545–1614.

42. Haberthür U, Caflisch A (2008) FACTS: Fast analytical continuum treatment of solvation. J Comput Chem 29(5):701–715.

