## Supplemental tables and figures for "Locating ligand binding sites in G-protein coupled receptors using combined information from docking and sequence conservation"

SUPPLEMENTARY INFORMATION

**Figure S1. Morphine docked to active conformations of a) β2 adrenergic and b) A2A adenosine receptors.** Docked with SwissDock to pdb 4ldo (β2 adrenergic) and 2ydo (A2A adenosine).

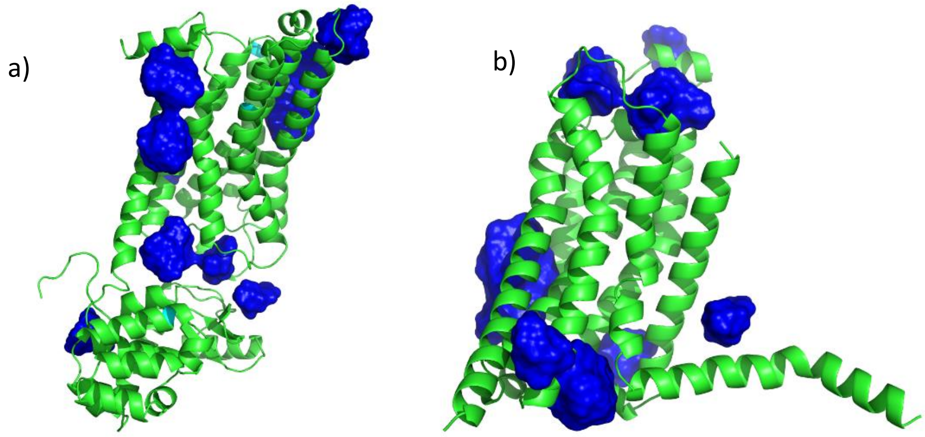

**Figure S2. Four experimentally verified GPER ligands**

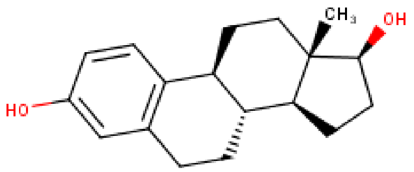

17β-estradiol

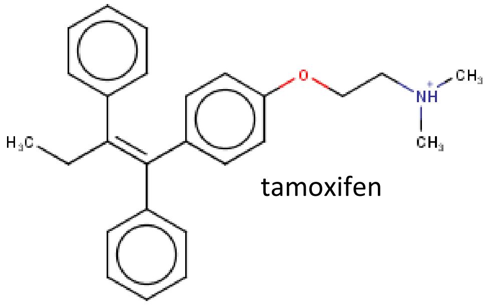

tamoxifen

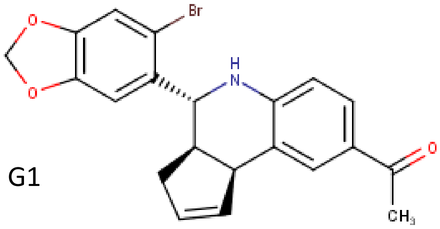

G1

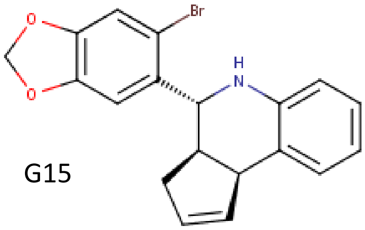

G15

**Figure S3. Overlay of GPER homology model with the crystal structure of the CCR5 chemokine receptor.** GPER is green. CCR5 receptor is blue. E2 poses are in blue. The top of the figure corresponds to the extracellular

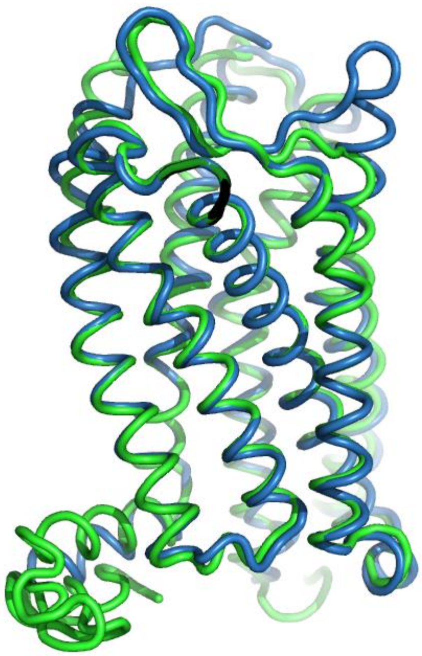

**Figure S4. Ramachandran plot of the GPER homology model.** Calculated by Chimera.

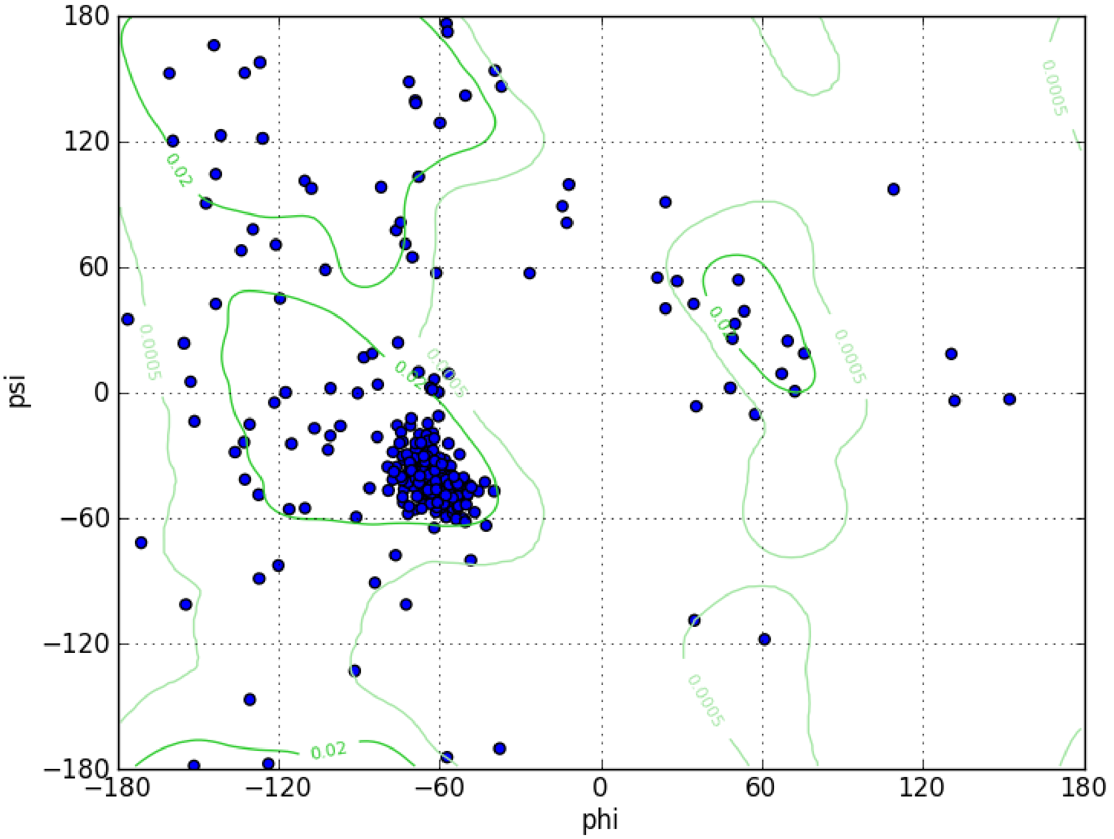

**Figure S5. ERRAT analysis of the GPER homology model.**

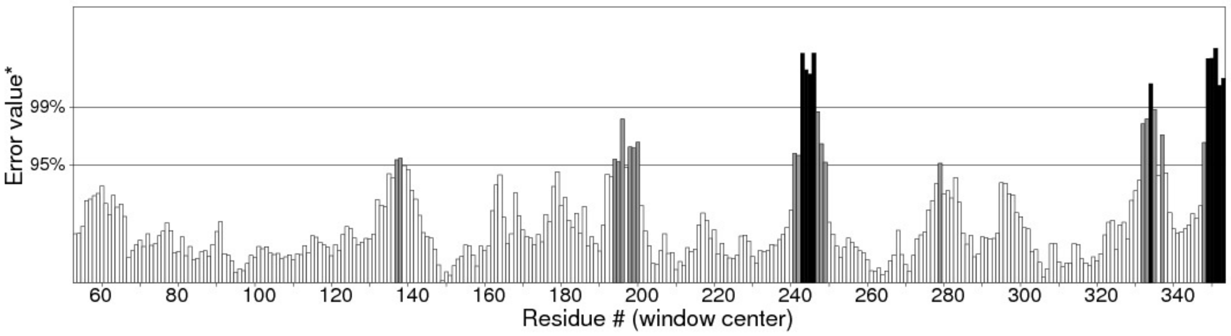

**Figure S6. E2 binding sites calculated by SwissDock.** E2 poses are in blue. The top of the figure corresponds to the extracellular face of GPER.

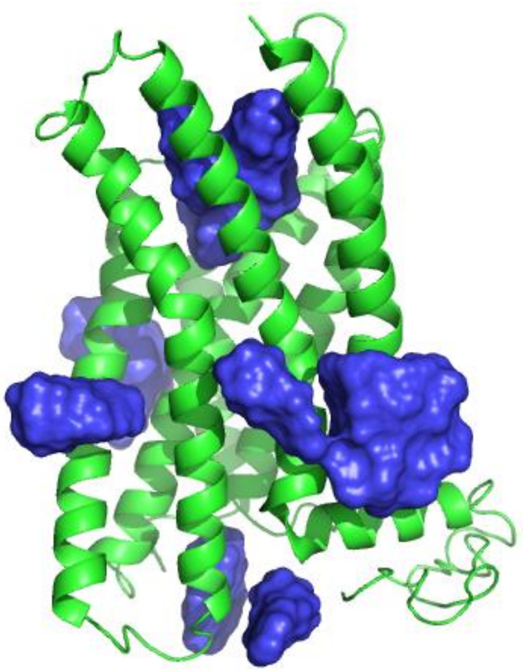

**Figure S7. Comparison of proposed ligand binding sites.** Comparison of the ConDock ligand binding site (red) with that proposed by Mendez-Luna *et al.* (blue). Residues found in both sites are colored violet.

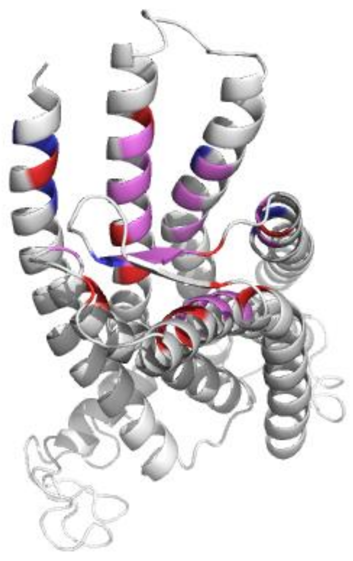

**Figure S8. ConSurf multiple sequence alignment for GPER.**

### ConSurf Color-Coded MSA

```
001 F7EQ49     MEVTSQARGMGLEMYPGTMQPAAPNTTSPELNLSHPLLGASLANGTGELS
002 O08878     MAATTPAQDVGVEIYLGPVWPAPSNSTPLALNLSLALREDAPGNLTGDLS
003 B3G515     MEEQ---TTNVIQIYVNG-------T--EQFNASF---DFNITDVKESTD
004 Q99527     MDVTSQARGVGLEMYPGTAQPAAPNTTSPELNLSHPLLGTALANGTGELS
005 B0F9W3     MEEQ---TTSLVWIYVNS-------T--EQLNTSY---EYNTTYLIEDSD
006 Q8BMP4     MDATTPAQTVGVEIYLGPVWPAPSNSTPLALNLSLALREDAPGNLTGDLS

001 F7EQ49     EHQQYVIGLFLSCLYTIFLFPIGFVGNILILVVNISFREKMTIPDLYFIN
002 O08878     EHQQYVIALFLSCLYTIFLFPIGFVGNILILVVNISFRERMTIPDLYFIN
003 B3G515     TYEFYIIGLFLSCLYTIFLFPIGFIGNILILVVNLNHRERMTIPDLYFVN
004 Q99527     EHQQYVIGLFLSCLYTIFLFPIGFVGNILILVVNISFREKMTIPDLYFIN
005 B0F9W3     KYQSYVIGLFLSCLYTILLFPIGFIGNILILVVNLNHRGKMAIPDLYFVN
006 Q8BMP4     EHQQYVIALFLSCLYTIFLFPIGFVGNILILVVNISFREKMTIPDLYFIN

001 F7EQ49     LAVADLILVADSLIEVFNLHEQYYDIAVLCTFMSLFLQVNMYSSVFFLTW
002 O08878     LAAADLILVADSLIEVFNLDEQYYDIAVLCTFMSLFLQINMYSSVFFLTW
003 B3G515     LAVADLILVADSLIEVFNLNEKYYDYAVLCTFMSLFLQVNMYSSIFFLTW
004 Q99527     LAVADLILVADSLIEVFNLHERYYDIAVLCTFMSLFLQVNMYSSVFFLTW
005 B0F9W3     LAVADLILVADSLIEVFNLNEKYYDYAVLCTFMSLFLQVNMYSSIFFLTW
006 Q8BMP4     LAAADLILVADSLIEVFNLDEQYYDIAVLCTFMSLFLQINMYSSVFFLTW

001 F7EQ49     MSFDRYIALARAMRCSLFRTKHHARLSCGLIWMASVSATLVPFTAVHLQH
002 O08878     MSFDRYLALAKAMRCGLFRTKHHARLSCGLIWMASVSATLVPFTAVHLRH
003 B3G515     MSFDRYVALTSSMSSSPLRTMQHAKLSCSLIWMASILATLLPFTIVQTQH
004 Q99527     MSFDRYIALARAMRCSLFRTKHHARLSCGLIWMASVSATLVPFTAVHLQH
005 B0F9W3     MSFDRYIALANSMSSSPLRTMQHAKLSCGLIWMASILATLLPFTIVQTQH
006 Q8BMP4     MSFDRYLALAKAMRCGLFRTKHHARLSCGLIWMASVSATLVPFTAVHLRH

001 F7EQ49     TDEACFCFADVREVQWLEVTLGFIVPFAIIGLCYSLIVRVLVRAHRHRGL
002 O08878     TEEACFCFADVREVQWLEVTLGFIVPFAIIGLCYSLIVRALIRAHRHRGL
003 B3G515     TGEVHFCFANVFEIQWLEVTIGFLIPFSIIGLCYSLIVRTLMRAQKHKGL
004 Q99527     TDEACFCFADVREVQWLEVTLGFIVPFAIIGLCYSLIVRVLVRAHRHRGL
005 B0F9W3     RGEVHFCFANVFEIQWLEVTIGFLVPFSIIGLCYSLIGRILMRSQKHRGL
006 Q8BMP4     TEEACFCFADVREVQWLEVTLGFIMPFAIIGLCYSLIVRALIRAHRHRGL

001 F7EQ49     RPRRQKALRMILAVVLVFFVCWLPENVFISVHLLQRTQPGAAPCKQSFRH
002 O08878     RPRRQKALRMIFAVVLVFFICWLPENVFISVHLLQWAQPGDTPCKQSFRH
003 B3G515     WPRRQKALRMIVVVVLVFFICWLPENVFISIQLLQGTADPSKRTDTTLWH
004 Q99527     RPRRQKALRMILAVVLVFFVCWLPENVFISVHLLQRTQPGAAPCKQSFRH
005 B0F9W3     WPRRQKALRMIVVVVLVFFICWLPENVFISIQLLQGTADPSQRTATTLRH
006 Q8BMP4     RPRRQKALRMIFAVVLVFFICWLPENVFISVHLLQWTQPGDTPCKQSFRH

001 F7EQ49     AHPLTGHIVNLAAFSNSCLNPLIYSFLGETFREKLRLYIEQKTNLPALNR
002 O08878     AYPLTGHIVNLAAFSNSCLSPLIYSFLGETFRDKLRLYVAQKTSLPALNR
003 B3G515     DYPLTGHIVNLAAFSNSCLNPIIYSFLGETFRDKLRLFIKRKASWSVVYR
004 Q99527     AHPLTGHIVNLAAFSNSCLNPLIYSFLGETFRDKLRLYIEQKTNLPALNR
005 B0F9W3     DYPLTGHIVNLAAFSNSCLNPIIYSFLGETFRDKLRLFIKQKASWSVVNR
006 Q8BMP4     AYPLTGHIVNLAAFSNSCLNPLIYSFLGETFRDKLRLYVEQKTSLPALNR
```

| | |
|---|---|
| 001 F7EQ49 | F C H A A L K A V I P D S T E Q S D V R F S S A V |
| 002 O08878 | F C H A T L K A V I P D S T E Q S D V K F S S A V |
| 003 B3G515 | F C N H T L D L Q I P V R S E S E V - - - - - - - |
| 004 Q99527 | F C H A A L K A V I P D S T E Q S D V R F S S A V |
| 005 B0F9W3 | F C H H G L D L H L P V R S E V S E V - - - - - - |
| 006 Q8BMP4 | F C H A T L K A V I P D S T E Q S E V R F S S A V |

| 1 | 2 | 3 | 4 | 5 | 6 | 7 | 8 | 9 |
|---|---|---|---|---|---|---|---|---|

Variable Average Conserved

X - Insufficient data - the calculation for this site was performed on less than 10% of the sequences.
